# *Arabidopsis thaliana* Trihelix Transcription factor AST1 mediates abiotic stress tolerance by binding to a novel AGAG-box and some GT motifs

**DOI:** 10.1101/121319

**Authors:** Hongyun Xu, Lin He, Yong Guo, Xinxin Shi, Dandan Zang, Hongyan Li, Wenhui Zhang, Yucheng Wang

## Abstract

Trihelix transcription factors are characterized by containing a conserved trihelix (helix-loop-helix-loop-helix) domain that bind to GT elements required for light response, play roles in light stress, and also in abiotic stress responses. However, only few of them have been functionally characterised. In the present study, we characterized the function of AST1 (Arabidopsis SIP1 clade Trihelix1) in response to abiotic stress. AST1 shows transcriptional activation activity, and its expression is induced by osmotic and salt stress. The genes regulated by AST1 were identified using qRT-PCR and transcriptome assays. A conserved sequence highly present in the promoters of genes regulated by AST1 was identified, which is bound by AST1, and termed AGAG-box with the sequence [A/G][G/A][A/T]GAGAG. Additionally, AST1 also binds to some GT motifs including GGTAATT, TACAGT, GGTAAAT and GGTAAA, but failed in binding to GTTAC and GGTTAA. Chromatin immunoprecipitation combined with qRT-PCR analysis suggested that AST1 binds to AGAG-box and/or some GT motifs to regulate the expression of stress tolerance genes, resulting in reduced reactive oxygen species, Na^+^ accumulation, stomatal apertures, lipid peroxidation, cell death and water loss rate, and increased proline content and reactive oxygen species scavenging capability. These physiological changes mediated by AST1 finally improve abiotic stress tolerance.

## Introduction

Trihelix transcription factors are characterized by a conserved trihelix (helix-loop-helix-loop-helix) domain that binds specifically to GT elements required for the light response, and are also termed GT factors (Zhou, 1999; Nagano *et al*., 2001). Compared with other transcription factor families, the trihelix family is relatively small, having 30 members in *A. thaliana* and 31 members in rice. According to the structure of the trihelical domain, the trihelix family is classified into five groups, GT-1, GT-2, SH4, GTγ and SIP1 (Kaplan-Levy *et al*., 2012; Qin *et al*., 2014).

The trihelix family binds to light-responsive GT elements in target promoters. These GT elements have different sequences, including GGTTAA, GGTAATT, GGTAAAT, GTTAC, TACAGT and GGTAAA, and are found in the promoters of light regulated genes, and are mainly involved in the light response (Green *et al*., 1987; Kay *et al*., 1989; O’Grady *et al*., 2001; Gao *et al*., 2009; Yoo *et al*., 2010). Moreover, GT elements are involved in biotic or abiotic stress responses, being found in many promoters of genes associated with drought, salt stress and pathogen infection (Buchel *et al*., 1996; Park *et al*., 2004; Yoo *et al*., 2010).

Trihelix transcription factors mainly respond to light stress and regulate the expression of light-responsive genes (Kaplan-Levy *et al*., 2012). Trihelix proteins are also involved in various developmental processes, including chloroplasts, embryonic development, seed germination and dormancy, stomatal aperture and the developments of trichomes and flowers (Willmann *et al*., 2011; Kaplan-Levy *et al*., 2014; O’Brien *et al*., 2015; Wan *et al*., 2015). Additionally, trihelix proteins are have roles in abiotic stresses, such as cold, oxygen, drought and salt stresses. For example, Arabidopsis GT-2 LIKE 1 (GTL1) is involved in plant water stress responses and drought tolerance, and *gtl1* mutations regulate stomatal density by reducing leaf transpiration to improve water use efficiency (Yoo *et al*., 2010). Poplar GTL1 has functions in water use efficiency and drought tolerance; when exposed to environmental stresses, PtaGTL1 induces Ca^2^+ signatures to modulate stomatal development and regulate plant water use efficiency (Weng *et al*., 2012). Arabidopsis GT-4 Trihelix can improve plant salt stress tolerance by regulating the expression of Cor15A to protect plants from the damage to the chloroplast membrane and enzymes caused by salt stress (Wang *et al*., 2014). *A. thaliana* AtGT2L and rice OsGTγ-1 were both induced by salt, drought, cold stress and abscisic acid (ABA) treatment (Fang *et al*., 2010; Xi *et al*., 2012). Although trihelix have roles in plants’ adaptation to various environmental stresses, their mechanisms of action in abiotic stress tolerance are largely unknown. For example, besides binding to GT motifs, whether they bind other cis-acting elements to regulate gene expression in response to abiotic stress, the identities of the target genes regulated by Trihelix and the physiological response mediated by Trihelix when exposed to abiotic stress remain to be revealed.

The function of Arabidopsis SIP1 clade Trihelix1 (*AST1*, At3g24860), which belongs to the trihelix subfamily of SIP1, has not been characterized. In this study, we characterized the function of AST1 in response to abiotic stress. Our study showed that AST1 plays an important role in plant salt and osmotic stresses, and we revealed the physiological responses mediated by AST1. Additionally, we identified a novel cis-acting element (the AGAG-Box) that is recognized by AST1. AST1 regulates stress-related genes by binding to the AGAG-Box and/or GT motifs to mediate salt and osmotic stress tolerance.

## Materials and methods

### Plant materials

*A. thaliana* ecotype Columbia was used in this study. The AST1 (AT3G24860) T-DNA insertion mutants, SALK_038594C, were obtained from the Arabidopsis Biological Resource Centre (ABRC). Three-week-old *A. thaliana* plants were watered with 150 mM NaCl or 200 mM mannitol for 3, 6, 12, 24 and 48 h, respectively. Roots and leaves were harvested for expression analysis, and plants treated with fresh water were also harvested at the corresponding time points as controls.

### Beta-glucuronidase (GUS) Staining and GUS Activity Quantification

The 1500 bp promoter of *AST1* together with the full 5′ UTR of *AST1* replaced the CaMV 35S promoter in vector PBI121 to drive *GUS* gene expression (ProAST1:GUS), and transformed into *A. thaliana* using the floral dip method (Clough *et al*., 1998). The T3 homozygous transgenic plants at different developmental stages were used for GUS staining and activity assays according to the methods described by Cheng *et al*. (2013) and Lu *et al*. (2007).

### Subcellular Localization assay

The coding sequence (CDS) of *AST1* was fused to the N-terminus of the green fluorescent protein (GFP) gene, under the control of CaMV 35S promoter (35S:AST1-GFP) and GFP under the control of 35S promoter was also generated (35S:GFP), and were transformed into *A. thaliana* plants. The root tips of 5-day-old transgenic seedlings were visualized using a fluorescence microscope Imager (Zeiss,Germany). The construct of 35S:AST1-GFP and 35S:GFP were also transformed separately into onion epidermal cells using the particle bombardment method and visualized using a confocal laser scanning microscopy LSM410 (Zeiss, Jena, Germany).

### Overexpression and knockout of AST1 in *A. thaliana*

The CDS of AST1 was inserted into pROK2 (Hilder *et al*., 1987) under the control of 35S promoter (35S:AST1), and were transformed into *A. thaliana*. Empty pROK2 was also transformed as the control. The expression of *AST1* in T3 homozygous transgenic lines or SALK_038594C plants was monitored by quantitative real-time reverse transcription PCR (qRT-PCR).

### Stress tolerance analysis

*A. thaliana* seeds were placed on 1/2 MS solid medium supplied with 150, 185 mM mannitol or 100, 125 mM NaCl for 10 days, and the proportion of seedlings survival rate was calculated. The 4-d-old seedlings grown on 1/2 MS solid medium were transferred to 1/2 MS medium supplied with 100 and 125 mM NaCl or 200 and 300 mM mannitol for 12 days, and the root lengths and fresh weights were measured. Three-week-old plants grown in the soil were treated separately with 150 mM NaCl or 200 mM mannitol for 10 days, and their fresh weights and chlorophyll contents were calculated; total chlorophyll contents were measured following the method of Gitelson *et al*. (2003).

### Detection of reactive oxygen species (ROS) and cell death

Three-week-old *A. thaliana* under normal conditions were watered with 150 mM NaCl or 200 mM mannitol for 24 h. To detect H_2_O_2_ and O_2_^.-^ content, leaves were infiltrated with nitroblue tetrazolium (NBT) or 3, 30-diaminobenzidine (DAB) solutions, as described by Zhang *et al*. (2011). For cell death determination, the detached leaves were incubated in Evans blue solution and stained according to Kim *et al*. (2003). For propidium iodide (PI) staining, 7-d-old seedlings in plates were treated with 150 mM NaCl or 200 mM mannitol for 24 h, and were used for PI staining according to Jones *et al*. (2016).

### Physiological analysis

Three-week-old *A. thaliana* plants under normal conditions were watered with 150 mM NaCl or 200 mM mannitol for 5 days, and were used for the following physiological measurements. Electrolyte leakage rate analysis was performed following the procedures described by Fan *et al*. (1997), and malonic dialdehyde (MDA) was determined according to the method of Madhava *et al*. (2000). Peroxidase (POD) and Superoxide Dismutase (SOD) were assayed as described previously (Han *et al*., 2008). The water loss rate was determined according to Hsieh *et al*. (2013). The proline content was determined as described by Han *et al*. (2014).

### Stomatal Aperture Analysis

Lower epidermal peels of 3-week-old plants leaves were stripped to float in a solution of 10 mM MES-KOH, pH 6.15, with 30 mM KCl, and were incubated under light for 2.5 h at 22°C to open the stomata. The leaves were then transferred to MES-KCl buffer, including 150 mM NaCl or 200 mM mannitol, for 3 h. Stomatal apertures were viewed using a light microscope (Olympus BX43, Japan) and measured by the software IMAGEJ 1.36b (http://brokensymmetry.com) (Watkins et al., 2014).

### Quantitative real-time reverse transcription PCR (qRT-PCR)

Total RNA was isolated using TRIzol reagent (Invitrogen). RNA (1 μg) was reverse transcribed into cDNA using oligo(dT) as primers, and diluted to 100 μl. For qRT-PCR, the reaction system (20 μl) included 10 μl of SYBR Green Realtime PCR Master Mix, 10 μM of forward or reverse primer and 2 μl cDNA dilution products. *ACT7* (*AT5G09810*) and *TUB2* (*AT5G62690*) were used as internal controls. All primers for qRT-PCR were shown in Table S1. The PCR was performed with an Opticon 2 System (Bio-Rad, Hercules, CA, USA) with following conditions: 94°C for 2 min; 45 cycles of 94°C for 30 s, 58°C for 30 s, 72°C for 40 s; and 79 °C for 1 s for plate reading. The relative expression levels were calculated using delta-delta Ct method (Livak and Schmittgen *et al*., 2001).

### Visualization and Measurement of Na^+^ and K^+^ contents

One week-old *A. thaliana* seedlings grown under normal conditions were treated with 150 mM NaCl for 24 h, and seedlings grown under normal conditions were used as controls. The plants were stained with 10 μM CoraNa-Green (Sigma, USA) for 2 h in the dark, then the root tips was visualized under an LSM710 microscope (Zeiss, Jena, Germany). After 150 mM NaCl or water treatment for 5 days, the roots and leaves were harvested for Na^+^ and K^+^ content analysis, which were performed as described preciously (Han *et al*., 2014).

### RNA-seq

Three-week-old *AST1* over-expressing plants and SALK plants were treated with 200 mM Mannitol for 24 h, and then the leaves were harvested for RNA-Seq. Statistical selection of differentially expressed genes between overexpression line 3 (OE3) and knockout line 2 (KO1.2) was based on a minimal 2.5 log2 fold change, together with a P-value ≤ 0.05 for the t-test, for three biological repetitions.

### MEME analysis

The promoter sequences (from −1 to −1000 bp) of the genes that are upregulated by AST1 were analyzed using the MEME program (http://meme-suite.org/tools/meme), with the same parameters used by Bailey *et al*. (2006).

### Yeast One-Hybrid (Y1H) Assays

The CDS of *AST1* was inserted into vector pGADT7-rec2 (Clontech) as the prey and one copy of each conserved sequence predicted by MEME was cloned into pHIS2 as baits (the primers were listed in Table S2). The positive clones were screened on SD/-Leu/-Trp (DDO) or SD/-His/-Leu/-Trp (TDO) medium supplied with 3-AT (3-Amino-1, 2, 4-triazole).

### Transient Expression Assay

The sequences that were confirmed to interact with AST1 by Y1H were cloned separately into a reformed pCAMBIA1301 vector (where 35S:hygromycion had been deleted, and a 46 bp minimal promoter was inserted between the BglII site and ATG of GUS) as reporter constructs (the primers were listed in Table S3). The 35S:AST1 was used as effector vector. The reporters and effector vector were co-transformed into tobacco by the transient transformation method (Zang *et al*., 2015), and 35S:LUC was cotransformed to normalize transformation efficiency. The GUS and LUC activities were determined as described previously (Lu *et al*., 2007).

### Electromobility shift assay (EMSA)

The CDS of AST1 was cloned into the pMAL-c5X vector between the BamHI and EcoRI enzyme digest sites and were induced to express by IPTG into *Escherichia coli* strain ER2523. Then the AST1 protein was extracted and purified following the Instruction Manual (NEB, pMAL^™^ Protein Fusion & Purification System). The probes were labeled with biotin using EMSA Probe Biotin Labeling Kit according to the manuals (Beyotime, China), and the unlabeled probe was used for the competitor. The EMSA was performed using Chemiluminescent EMSA kit (Beyotime, China). The primers used for EMSA were listed in Table S4

### ChIP Assays

Three-week-old *A. thaliana* expressing the AST1-GFP fusion gene were used for ChIP analysis. The plants were treated with 150 mM NaCl or 200 mM Mannitol for 24 h, and then harvested for the ChIP assays. ChIP experiments were performed as described by Haring *et al*. (2007). The cross-linked chromatin was sonicated and incubated with an anti-GFP antibody (Beyotime, China) (ChIP+), and the chromatin incubated with a rabbit anti-haemagglutinin (HA) antibody was used as the negative control (ChIP-). The DNA was detected by qPCR with the CDS of *Actin2* (*At3G18780*) as an internal control. The primers used for ChIP were listed in Table S5.

### Accession Numbers

Sequence data from this article can be found in The Arabidopsis Information Resource (http://www.arabidopsis.org/) under the following accession numbers: SOD2(AT2G28190), FSD1(AT4G25100), SOD(AT5G11000), SOD(AT3G10920), PER4(AT1G14540), POD(AT1G24110), POD(AT1G71695), PRX37(AT4G08770), PRX72(AT5G66390), POD(AT5G58400), ATMYB61 (AT1G09540), P5CS1 (AT2G39880), P5CS2(AT3G55610), PRODH (AT4G34590), P5CDH (AT5G62520), HKT1 (AT4G10310), NHX2(AT3G05030), NHX3(AT5G55470), NHX6(AT1G79610), SOS2(AT5G35410), SOS3 (AT5G24270), LEA3(AT1G02820), LEA7(AT1G52690), COR15(AT2G42520), LEA14(AT1G01470), ATCOR47(AT1G20440), ERD10(AT1G20450), ABR (AT3G02480), LSU1 (AT3G49580), SAUR16 (AT4G38860).

## Results

### Spatial and temporal expression profiles of *AST1*

GUS staining was performed on the transgenic *A. thaliana* plant expressing ProAST1:GUS to determine the expression profile of *AST1. AST1* was expressed at each studied developmental stage and in different tissues. The expression of *AST1* increased from 5-d- to 20-d-old seedlings, but reduced in plants older than 20 d (Figure 1A, 1-6), displaying a temporal expression pattern. *AST1* was highly expressed in leaves, stems and anthers compared with roots and siliques (Figure 1A, 7-11). Consistently, qRT-PCR showed that *AST1* was highly expressed in stems, leaves and flowers, but had relative lower expression levels in roots and siliques (Figure 1B). Interestingly, although *AST1* was expressed in leaves, it had relative higher expression in guard cells, root and leaf vascular systems (Figure 1A8, 12-13).

**Figure 1.**
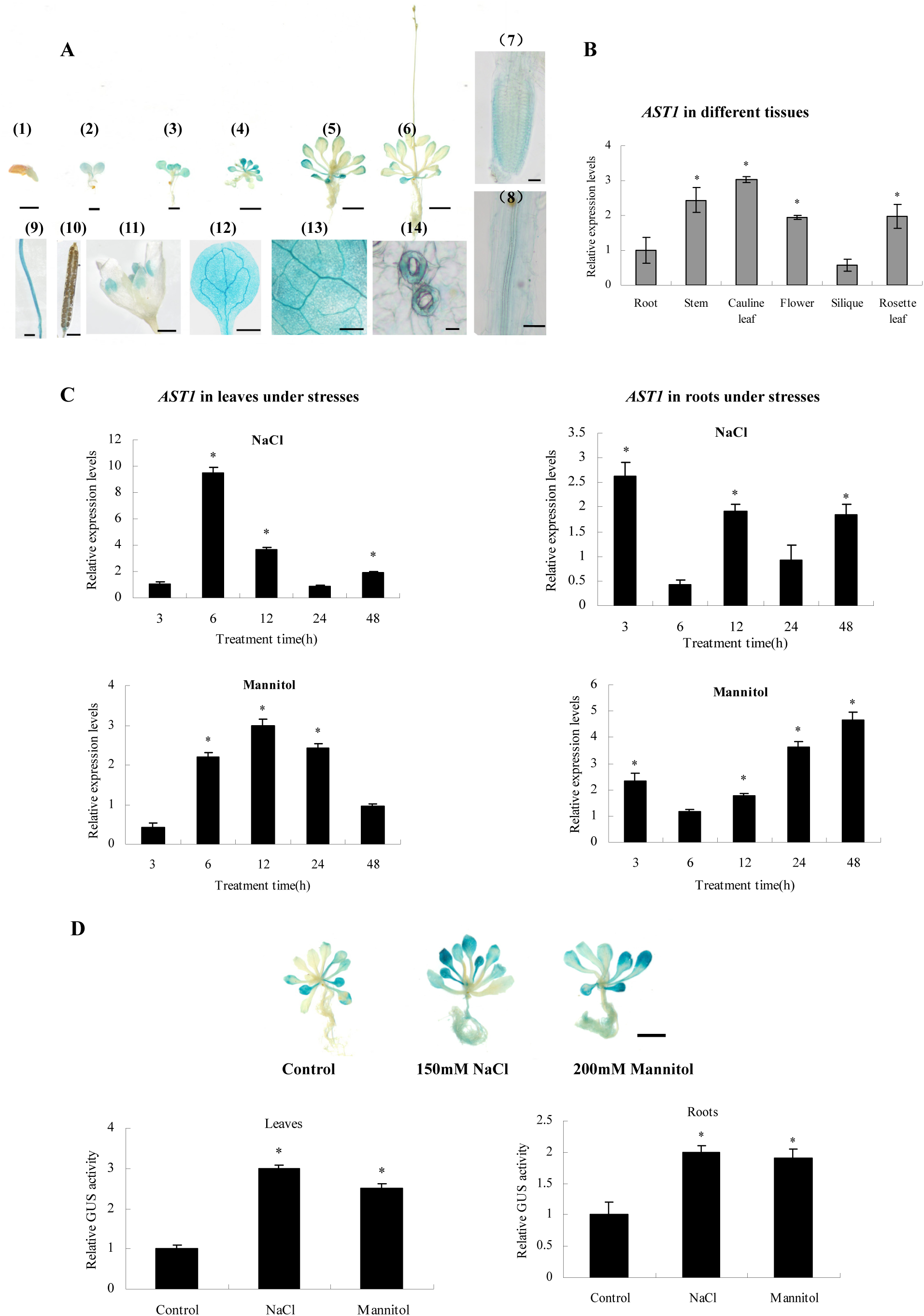
Expression profiles of AST1. **(A)** GUS staining analysis of ProAST1:GUS transgenic plants. (1−6) 2-, 5-, 10-, 15-, 20- and 30-d-old-seedling, (7) Root, (8) root vascular tissue, (9) Stem, (10) Silique, (11) Flower, (12) Rosette leave, (13) Leaf vascular tissue, (14) Guard cells. Bars: (1−3, 9, 10, 12,) 1 mm, (4−6) 1 cm, (7 and 8) 25 μm, (11) 250 μm, (13) 50 μm, (14) 10 μm. **(B)** The expression of *AST1* in different tissues of WT *A. thaliana* using qRT-PCR. Tissues from four-week-old plants were used for analysis. The expression level in roots was set as 1 to normalize the expression in other tissues. Asterisk (*) indicates significant difference compared with the roots (P < 0.05). **(C)** The expression of *AST1* in response to abiotic stresses. The expression level in the samples treated with fresh water harvested at each time point were as the controls, and was set as 1 to normalize the expression at the corresponding time point. Three biological replications were conducted. The error bars represent the standard deviation (S.D.). Asterisk (*) indicates significant difference between treatments and controls (P < 0.05). **(D)** GUS staining of ProAST1:GUS transgenic plants under abiotic stress conditions. *A. thaliana* plants containing ProAST1:GUS grown in 1/2 MS medium were treated with NaCl or Mannitol for 12 h. At least 10 seedlings were included in each experiment, and three biological replications were performed. The GUS activity in control sample (no stress) was set as 1 to normalize the activity under stress conditions. Bars: 1cm. Three biological replications were performed. Asterisk (*) indicates a significant difference between treatments and controls (P < 0.05).

Under NaCl stress conditions, in leaves, the expression of *AST1* was highly induced at 6 to 12 h, but continually decreased after 6 h of stress (Figure 1C). In roots, *AST1* was highly induced by stress for 3, 12 and 48 h, downregulated at 6 h, and recovered at 24 h under NaCl stress conditions. Under mannitol stress conditions, in leaves, the expression of *AST1* was downregulated at 3 h, but increased continually from 6 to 12 h, reaching its expression peak at 12 h, after which it decreased continually (Figure 1C). In roots, *AST1* was slightly induced by stress for 3 to 12 h, highly induced at 24 and 48 h, and reached its expression peak at mannitol stress for 48 h (Figure 1C). Consistently, determination of GUS activity in Arabidopsis plant expressing ProAST1:GUS also confirmed that the expression of *AST1* was significantly induced in leaves and roots after exposed to mannitol or NaCl for 12 h (Figure 1D). These results suggested that the expression of *AST1* responded to salt and mannitol stress, and might play a role in salt and osmotic stress tolerance.

### Subcellular localization of AST1

The results showed that the GFP signal was detected in the whole cells of root tips or root elongation zone in *A. thaliana* plants expressing 35S:GFP (Figure S1A). However, the GFP signal was only detected in the nucleus of the root tips to the root hair zone in *A. thaliana* expressing AST1-GFP (Figure S1A). Additionally, transient transformation of onion epidermal cells also indicated that AST1 was localized in the nucleus (Figure S1B). Taken together, these results indicated that AST1 was target to the nucleus.

### Generation of overexpression or knockout plants for *AST1*

The T3 generation of *A. thaliana* plants overexpressing *AST1* (OE) and the *AST1* mutant plants (SALK_038594C) (KO plants) were generated, and the T-DNA sequence was inserted at the position that was at the 388 bp down-stream of the ATG. The qRT-PCR results showed that the expression of *AST1* was significantly increased in the OE plants and highly decreased in the KO plants (Figure S2), indicating that *AST1* had been successfully overexpressed and knocked-out, respectively, and that these plants were suitable for gain and loss-of-function analysis. Three AST1-overexpressing lines (OE1, OE2 and OE3) that had relative high *AST1* expression and three homozygous mutant plants (KO1.1, KO1.2 and KO1.3) that had the lowest *AST1* expression were selected for further study. Wild-type plants (WT) and WT plants transformed with the empty pROK2 vector (35S) were used as the controls.

### AST1 improves Drought and Salt tolerance

Under normal conditions, there was no difference in seedling survival rates among all the studied plants (Figure 2A). Under salt or osmotic stress conditions, compared with the WT, all OE plants showed significantly higher seedling survival rates, all KO plants showed significant lower seedling survival rates, and 35S plants showed similar seedling survival rates (Figure 2A). Root length and fresh weight were analyzed to determine stress tolerance. There was no difference in growth phenotype, root elongation and fresh weights among all the studied lines under normal conditions (Figure S3). Under stress conditions, compared with WT plants, the root elongation and fresh weights of KO plants were significantly reduced, but all the OE plants showed significantly increased root elongation and fresh weights; the 35S plants were similar to the WT (Figure S3). Stress tolerance was further studied in seedlings grown in soil. There was no difference in growth phenotype, fresh weights, and chlorophyll content among the studied plants under normal conditions (Figure 2B). Under salt or osmotic stress conditions, compared with the WT and 35S plants, all the OE plants showed increased chlorophyll content and fresh weights, and the KO lines displayed decreased fresh weights and chlorophyll contents. These results suggested that overexpression or knockout of *AST1* didn’t affect the growth and phenotype of plants. However, *AST1* could regulate salt and osmotic stress tolerance positively.

**Figure 2.**
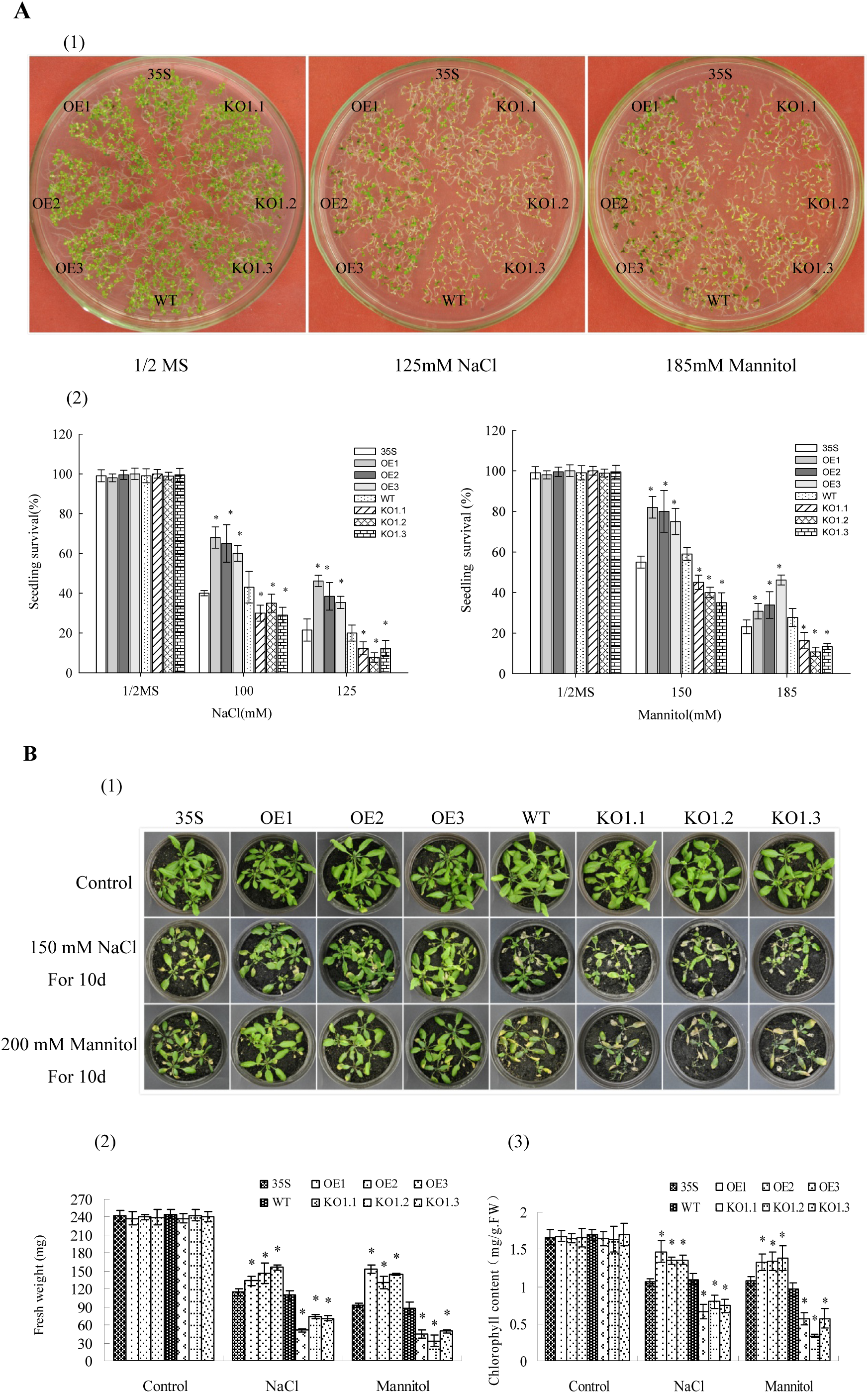
Abiotic Stress tolerance analysis of AST1. **(A)** Analysis of seed germination phenotypes under salt and osmotic conditions. (1) Germination phenotype. *A. thaliana* grown in 1/2 MS medium were treated with NaCl or Mannitol for 10 d. 35S: *A. thaliana* transformed with empty pROK2 (35S), OE: transgenic plants overexpressing AST1; WT: Wild Type; KO: *A. thaliana* mutant plants with knockout of AST1. The photographs showed representative seedlings. (2) Seed germination assay. The survival rates under NaCl (100 or 125 mM) or Mannitol (150 or 185mM) were calculated. *A. thaliana* plants grown in 1/2 MS medium were used as the control. Data are means ± SD from three independent experiments. Asterisk (*) indicates significant (t test, P < 0.05) difference compared with WT. **(B)** Stress tolerance analysis on seedlings grown in soil. (1) Three-week-old *A. thaliana* plants grown in soil were watered with 150 mM NaCl or 200 mM Mannitol for 10 d, well watered plants were used as the control. (1) The growth of *A. thaliana* plants under salt or osmotic stress for 10 d. (2, 3) Measurement of fresh weight and chlorophyll content. Bars indicate the mean ±standard deviation (SD) for each set of three independent experiments (n=30). (*P < 0.05). Significant difference compared with WT.

### Stomatal aperture and water loss rate analysis

As AST1 is highly expressed in guard cells (Figure 1A), we studied whether it played a role in controlling stomatal apertures. Under normal conditions, all the lines had similar stomatal apertures and width/length ratios (Figure 3A). When exposed to salt and osmotic stress, the WT and 35S plants had similar stomatal apertures and width/length ratios. Compared with the WT plants, the OE lines displayed decreased stomatal apertures and lower width/length ratios, and the KO plants showed increased stomatal apertures and higher width/length ratios (Figure 3A). Protein AtMYB61 was found to control the stomatal aperture ( Liang *et al*., 2005); therefore, we further studied whether AST1 could regulate *AtMYB61* expression. The expression of *AtMYB61* was significantly increased in the OE plants compared with the WT and 35S plants, and was significantly decreased in the KO plants (Figure 3B). The stomatal aperture is closely related with the water loss rate; therefore, we further studied the water loss rates under dehydration conditions. WT and 35S lines had similar water loss rates; however, the KO plants exhibited increased water loss rates, and the OE plants displayed decreased water loss rates compared the WT plants (Figure 3C). These results together indicated that AST1 regulates *AtMYB61* expression positively to control stomatal aperture, resulting in a reduced water loss rate.

**Figure 3.**
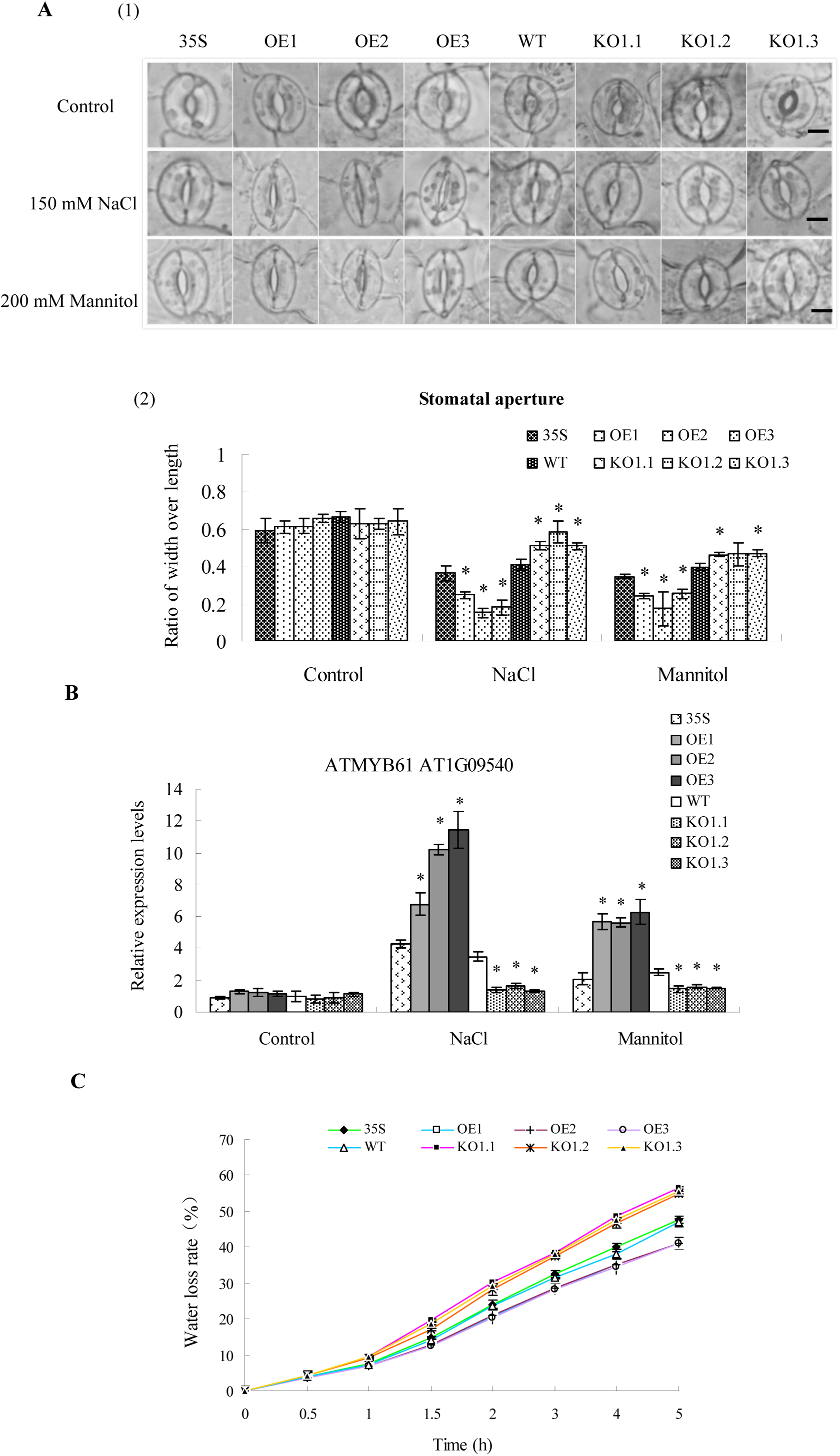
Comparison of Stomatal closure and water loss rates. **(A)** Stomatal closure assay. (1) The stomatal aperture under normal, salt and osmotic stress conditions. Stomata were pre-opened under light and then incubated in the solution of 150 mM NaCl or 200 mM Mannitol for 2.5 h under light. Water-mediated stomatal closure was used as a control. (2) Measurement of stomatal aperture. Values are mean ratios of width to length. Error bars represent standard errors of three independent experiments (n=30−50). Bars: 10 um. Asterisk (*) indicates a significant difference at P<0.05 compared with the WT. **(B)** Analysis of the expression of stomatal aperture-related gene *AtMYB61* (*AT1G09540*). The plants were treated with water (Control), 150 mM NaCl or 200 mM Mannitol, and the expression level of *AtMYB61* in WT plants under normal conditions was used to normalize all other expressions. Data are means ±SD from three independent experiments. Asterisk (*) indicates a significant difference at P<0.05 compared with the WT. **(C)** Analysis of water loss rates. Leaves from three-week-old plants were harvested for transpiration at room temperature. Values are means of the percentage of leaf water loss ± SD (n=30). Three independent experiments were performed.

### Determination of Na^+^ and K^+^ contents

The accumulation of Na^+^ in root tips was visualized by CoroNa-Green, a sodium-specific fluorophore. Under normal conditions, there was no substantial difference in Na^+^ accumulation among the studied plants. However, under salt stress conditions, KO plants displayed substantially stronger fluorescence than the WT and 35S plants, and the OE plants showed the weakest fluorescence (Figure 4A), indicating that Na^+^ was highly accumulated in the KO plants, but was accumulated lowly in the OE plants compared with in WT plants. Na^+^ and K^+^ contents were further determined using a Flame spectrophotometer. All the studied lines had generally similar Na^+^ and K^+^ contents under normal conditions. Under NaCl stress condition, Na^+^ was increased and K^+^ was decreased in all plants. However, in both the leaves and roots, the Na^+^ content was highly accumulated in KO plants, followed by the WT and 35S plants; the OE plants had lowest Na^+^ level (Figure 4B), which was consistent with CoroNa-Green staining. Meanwhile, the OE plants had higher K^+^ levels, and KO plants had lower K^+^ level compared with those in the WT and 35S plants in both leaves and roots (Figure 4B). The K^+^/Na^+^ ratios were similar in the leaves and roots of all plants under normal conditions. Under salt stress conditions, the OE plants had the highest K^+^/Na^+^ ratio, followed by the WT and 35S lines, and the KO plants had the lowest K^+^/Na^+^ ratio (Figure 4B). We further examined the expression of genes related to Na^+^ or K^+^ transport, including those encoding 1 sodium transporter (*HKT1*), and three Na^+^ (K^+^)/H^+^ transport proteins (*NHX2, NHX3, NHX6*), two salt overly sensitive (SOS) family proteins (*SOS2* and *SOS3*), which control plant K^+^ and Na^+^ nutrition. The results showed that *HKT1* had its highest expression in KO plants, followed by that in the WT and 35S plants, and was lowest in the OE plants (Figure 4C). Conversely, *NHX2, NHX3, NHX6* and *SOS2* showed their highest expression levels in the OE plants, followed by the WT and 35S plants, and showed their lowest expression levels in the KO plants. The expression of *SOS3* was not significantly different among the studied lines (Figure 4C).

**Figure 4.**
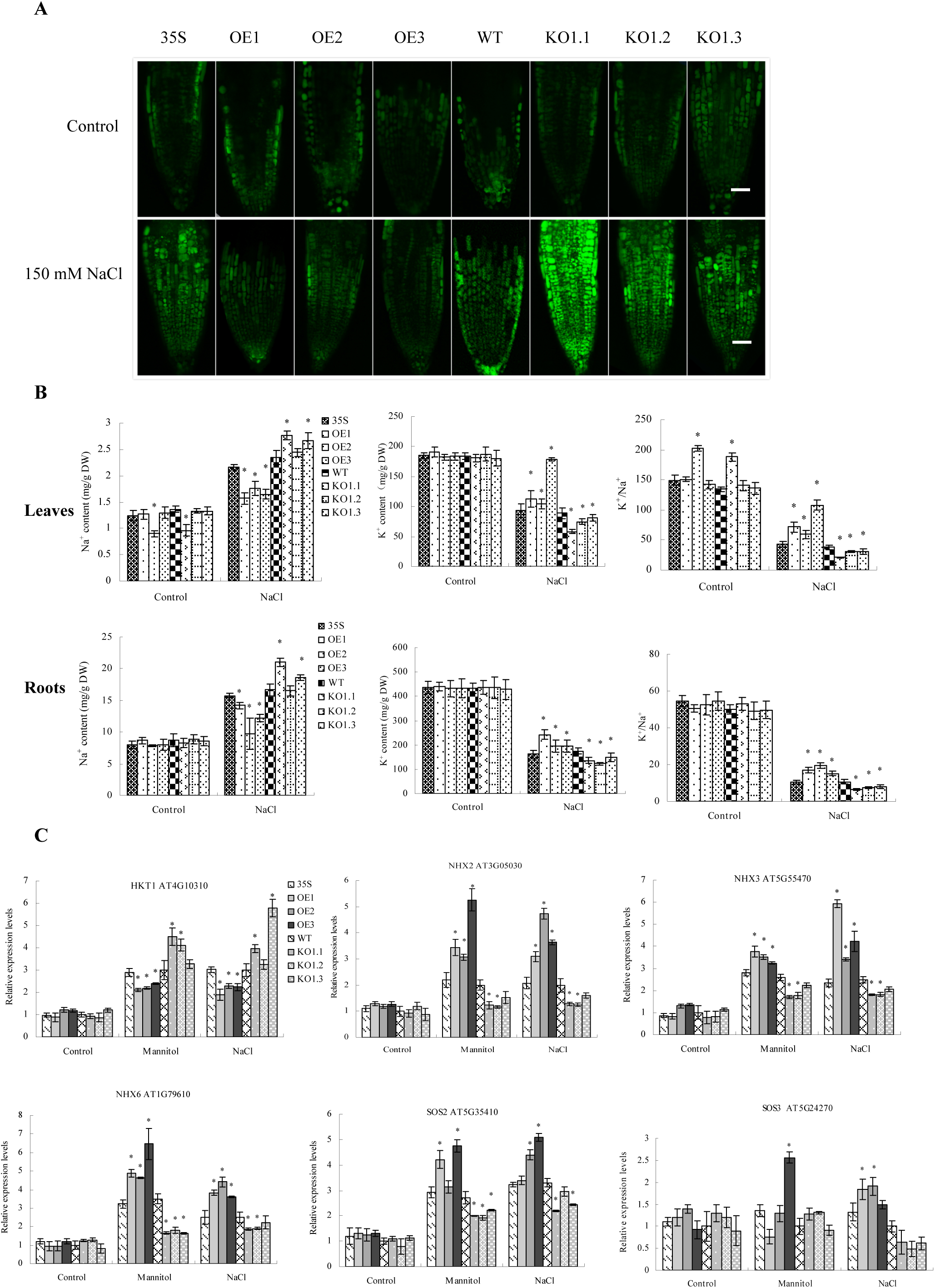
Analysis of Na^+^ and K^+^ contents. **(A)** Image of Na^+^ distribution in root tips. Five-d-old plants were treated with water (control) and 150 mM NaCl, respectively, for 24 h for staining with CoroNa-Green. The roots of thirty seedlings were used for each type of plant, and some roots were randomly selected to be photographed. **(B)** Measurement of Na^+^ and K^+^ contents in leaves and roots. Na^+^ and K^+^ were measured from 3-week -old plants of 150 mM NaCl treatment, and then K^+^/ Na^+^ ratio were respectively calculated. Results are presented as means and standard errors from three independent biological replicates. **(C)** The relative expression of genes involved in Na^+^ or K^+^ transporting. The expression of each gene in WT plants under normal conditions was set as 1 to normalize its expression in different lines under different conditions. Data are means ± SD from three independent biological replicates. Asterisk (*) indicates a significant difference at P<0.05 compared with the WT.

### Analysis of proline metabolism

Proline is an important osmotic adjustment substance and also plays a role in ROS scavenging; therefore we measured the proline contents in the plant lines. The results showed that all the lines had similar proline contents under normal conditions. However, when exposed to salt or osmotic stress, the OE plants had highest proline level, followed by the WT and 35S plants, and the KO plants had the lowest proline contents (Figure 5A). We further investigated the genes involved in proline metabolism, including two proline biosynthesis genes, D(1)-pyrroline-5-carboxylate synthetase (*P5CS*) gene, *p5CS1* and *p5CS2;* two proline degradation genes, D(1)-pyrroline-5-carboxylate dehydrogenase (*P5CDH*) and proline dehydrogenase (*PRODH*). When exposed to salt or osmotic stress conditions, the expression of both *p5CS1* and *p5CS2* were increased in the OE lines and decreased in the KO lines, compared with the WT and 35S lines. Conversely, *P5CDH* and *PRODH* showed the highest expression levels in the KO plants, followed by the WT and 35S lines, and the lowest level in the OE plants (Figure 5B). These results indicated that AST1 could increase proline content by affecting the expression of proline metabolism genes.

**Figure 5.**
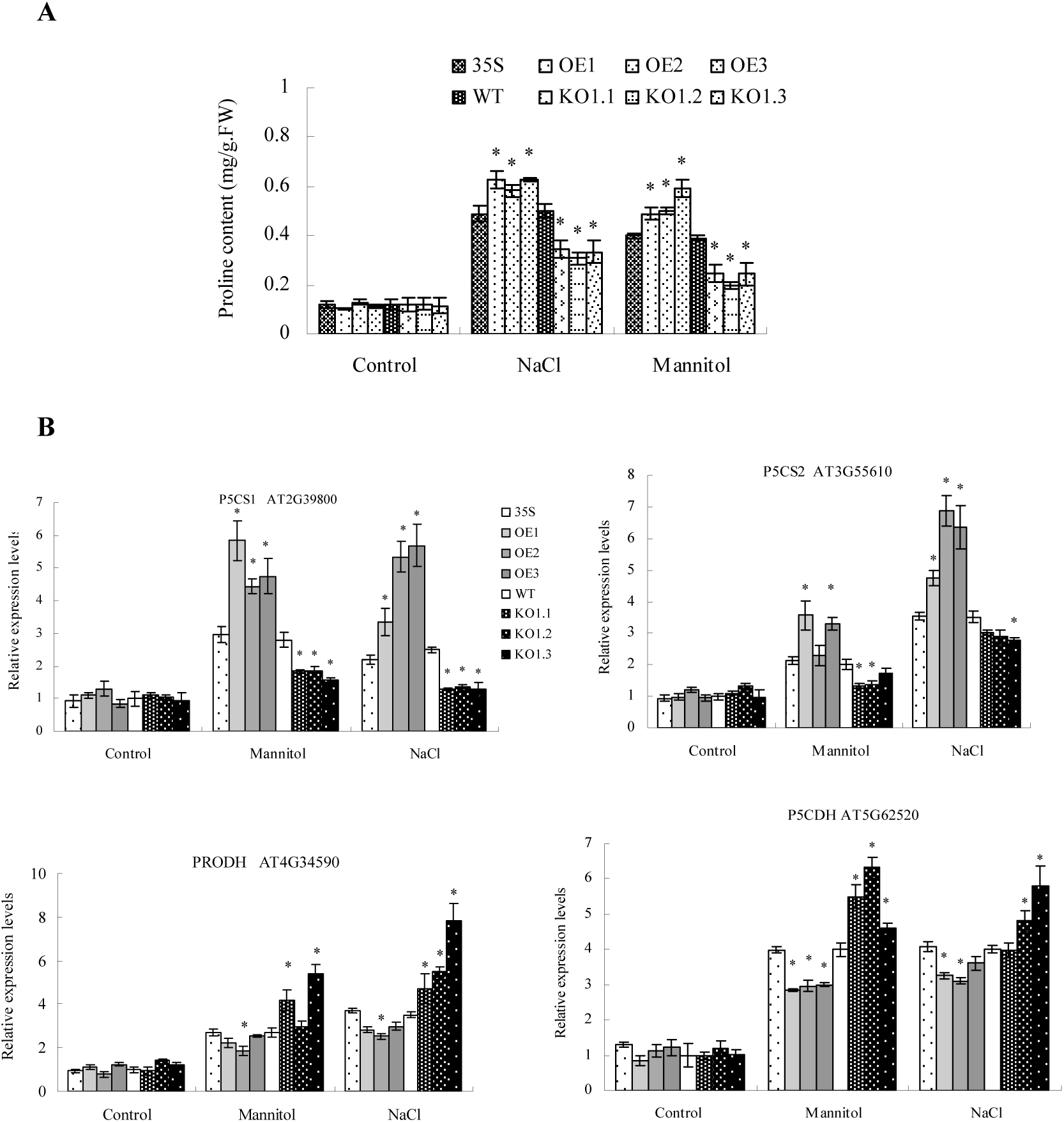
The regulation of proline metabolism genes by AST1. (A) Proline content assay. Values represent the average of three biological replicates. Significant differences from WT are indicated. (B) The transcripts level of proline metabolism genes. The expression of each gene in WT plants under normal condition was set as 1 to normalize its expression in different lines under different conditions. Asterisk (*) indicates a significant difference at P<0.05 compared with the WT.

### Cell death and MDA content analysis

Evans blue and PI fluorescence staining were used to detect cell death in leaves and roots, respectively. There was no difference in Evans blue and PI staining among all the plants under normal conditions. Under NaCl and mannitol conditions, compared with the WT and 35S plants (they have similar cell death rates according to the staining), both Evans blue and PI staining showed that cell death was substantially decreased in OE plants. By contrast, KO plants displayed increased cell death (Figure S4A, B). To measure cell death quantitatively, the electrolytic leakage rates were determined. All the studied lines shared similar electrolytic leakage rates under normal conditions. Under salt or osmotic stress, the WT and 35S plants shared similar electrolytic leakage rates; however, compared with the WT and 35S plants, all KO and OE plants showed increased and decreased electrolytic leakage rates, respectively (Figure S4C), which was consistent with the results from Evans blue and PI staining.

Malonic dialdehyde (MDA) contents were measured to evaluate the level of membrane lipid peroxidation. Under normal conditions, all the plants had similar MDA levels. Under salt or osmotic stress conditions, the KO plants had the highest MDA content, followed by the WT and 35S plants (they shared similar MDA level), and the OE plants showed the lowest MDA contents (Figure S4D). These results indicated that expression of *AST1* could reduce membrane lipid peroxidation under abiotic stress conditions.

### ROS scavenging assay

We first studied the contents of O_2_^.-^ and H_2_O_2_ by nitroblue tetrazolium (NBT) and 3, 30-diaminobenzidine (DAB) *in situ* staining, respectively, and a deeper the blue or brown color indicated the accumulation of O_2_^.-^ and H_2_O_2_, respectively. There was no observable difference in NBT and DAB staining among the WT, OE, 35S and KO plants under the normal conditions (Figure S5A). When exposed to NaCl or mannitol, the WT and 35S plants had similar O_2_^.-^ and H_2_O_2_ levels; compared with them, OE plants displayed substantially reduced O_2_^.-^ and H_2_O_2_ accumulation, and all KO plants showed increased O_2_^.-^ and H_2_O_2_ accumulation.

The reactive oxygen species (ROS) levels were altered; therefore, we further studied whether this was caused by altered ROS scavenging capability. Peroxidase (POD) and Superoxide Dismutase (SOD) activities were measured. Under normal conditions, there was no difference in SOD and POD activities among all the plants. However, under NaCl or mannitol conditions, the activities of SOD and POD in the OE plants were the highest, followed by the WT and 35S plants, and the KO plants had the lowest SOD and POD activities (Figure S5B). The expression of the *SOD* and *POD* genes were further studied, and the genes that have known SOD or POD activity were selected for study. Under salt and mannitol conditions, the expression levels of all the *POD* and *SOD* genes (except for *ATSOD1*) in OE plants were the highest, followed by the WT and 35S, and the KO plants had the lowest expression levels (Figure S5C). This result indicated that AST1 could induce the expression of *SOD* and *POD* genes to elevate the SOD and POD activities when exposed to salt and osmotic stress.

### AST1 induced the expression of *LEA* family genes in response to salt and drought stresses

Seven *LEA* (late embryogenesis abundant) family genes that had been reported to be involved in abiotic stress tolerance were studied. Under normal conditions, there was no difference in expression levels among the plants (Figure S6). When exposed to NaCl or Mannitol, except for the *ABA-RESPONSE PROTEIN* (*ABR*) gene, all the studied *LEA* family genes displayed their highest expression levels in the OE plants, followed by the WT and 35S plants, and showed their lowest expression levels in the KO plants (Figure S6). These results showed that *AST1* could induce certain *LEA* family genes to improve abiotic stress tolerance.

### RNA-Seq analysis

A transcriptomic analysis was carried out to identify differentially expressed genes (DEGs) between the OE3 and KO1.2 lines. In total, 144 DEGs (fold change < 2 and false discovery rate (FDR) < 0.05) were identified, among which 65 genes were upregulated and 77 genes were downregulated. These DEGs were listed in Table S6 and the hierarchical clustering analysis was shown in Figure S7. Gene ontology (GO) analysis showed that these DEGs were mainly involved in signaling, immune system process, reproduction and cell killing in biological process (Figure S8A). Kyoto encyclopedia of genes and genomes (KEGG) analysis showed that the DEGs were mainly associated with plant hormone signal transduction and plant-pathogen interaction pathways (Figure S8B). These results indicated that AST1 played a key role in regulating these pathways.

### A novel motif recognized by AST1

To study the motif mainly bound by AST1 in regulating gene expression when exposed to abiotic stress, the MEME motif discovery tool (http://meme-suite.org) was used. As shown in Figure 7, AST1 could bind to GT2, GT3, GT4 and GT5 to active gene expression, suggesting that AST1 should play an expressional activation role. Therefore, the promoters of 54 genes that were highly upregulated by AST1 according to the qRT-PCR and RNA-Seq analyses were employed for further study. The MEME results showed that there was a 12 base conserved sequence present in most of the studied promoters (Figure 6A).

**Figure 6.**
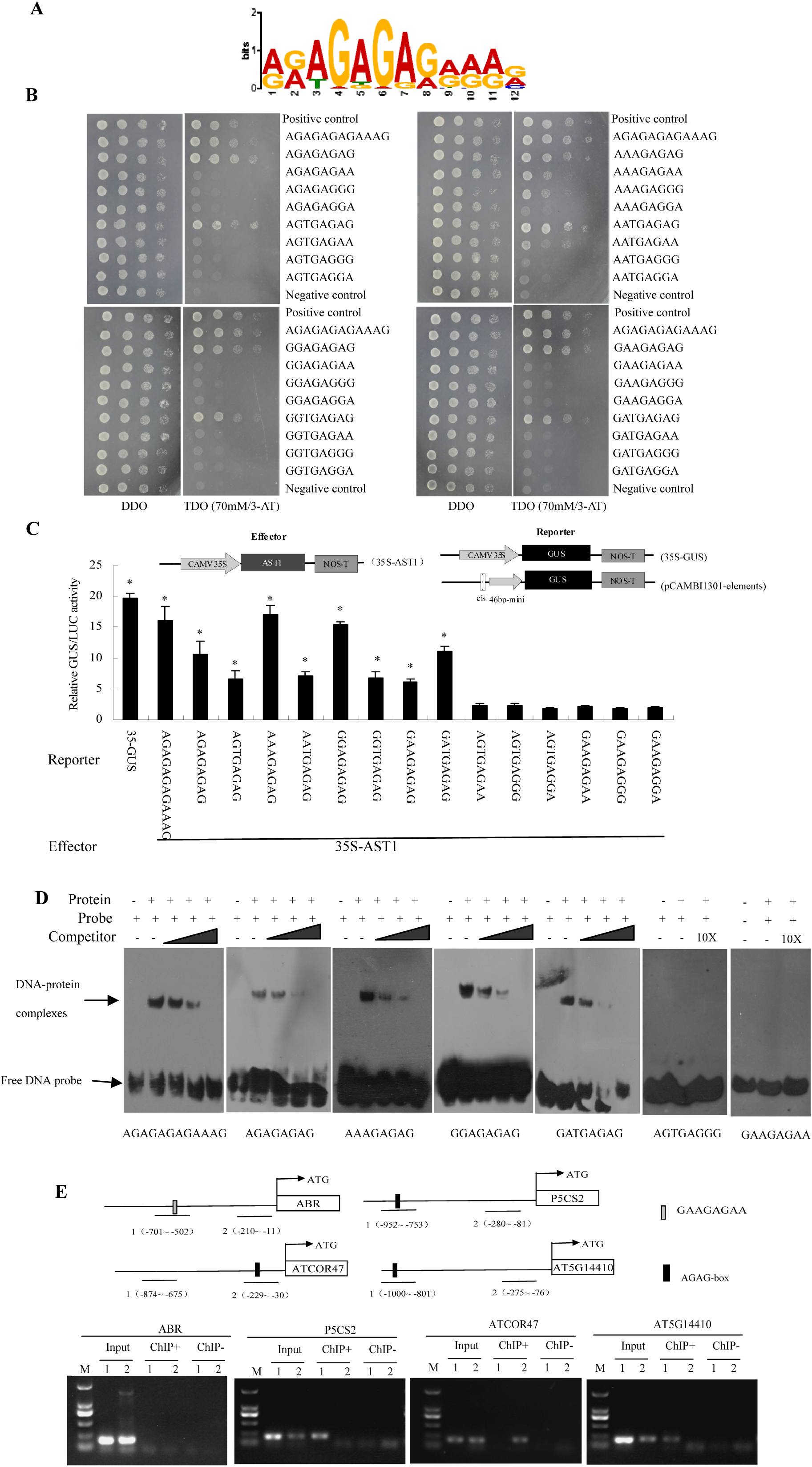
Identification of AGAG-box recognized by AST1. **(A)**MEME analysis of the conserved sequence present in the promoters of genes regulated by AST1. **(B)**Y1H assay of the interaction of AST1 with the full or the core conserved sequences. The 12 bp conserved sequence or the 1st to 8th base of conserved sequences (32 types in total) were tested for their interaction with AST1 using Y1H. **(C)** Determination of the interaction between AST1 and AGAG-box in tobacco plants. The studied sequences were fused separately with the 46-bp minimal promoter to drive a *GUS* gene as reporters, and were then co-transformed with 35S:AST1 and 35S:LUC into tobacco plants. Diagrams of the reporter and effector vectors were shown. Data are means ± SD from three independent biological replicates. Asterisk (*) indicates a significant difference at P<0.05 compared with the sequence “GAAGAGGA” **(D)** EMSA was carried out with AST1 protein and five type sequences of AGAG-box as probes. Competition for the labeled sequences was tested by adding 10-, 30- and 100-fold excess of unlabeled probes. The free probes and DNA-AST1 complexes were indicated with arrows. **(E)** ChIP analysis of the binding of AST1 to the AGAG-box in *A. thaliana*. The gene promoters that contained only one AGAG-box and did not contain any GT motifs bound by AST1 were used. Schematic diagram showing the positions of the AGAG-box in the promoters.

**Figure 7.**
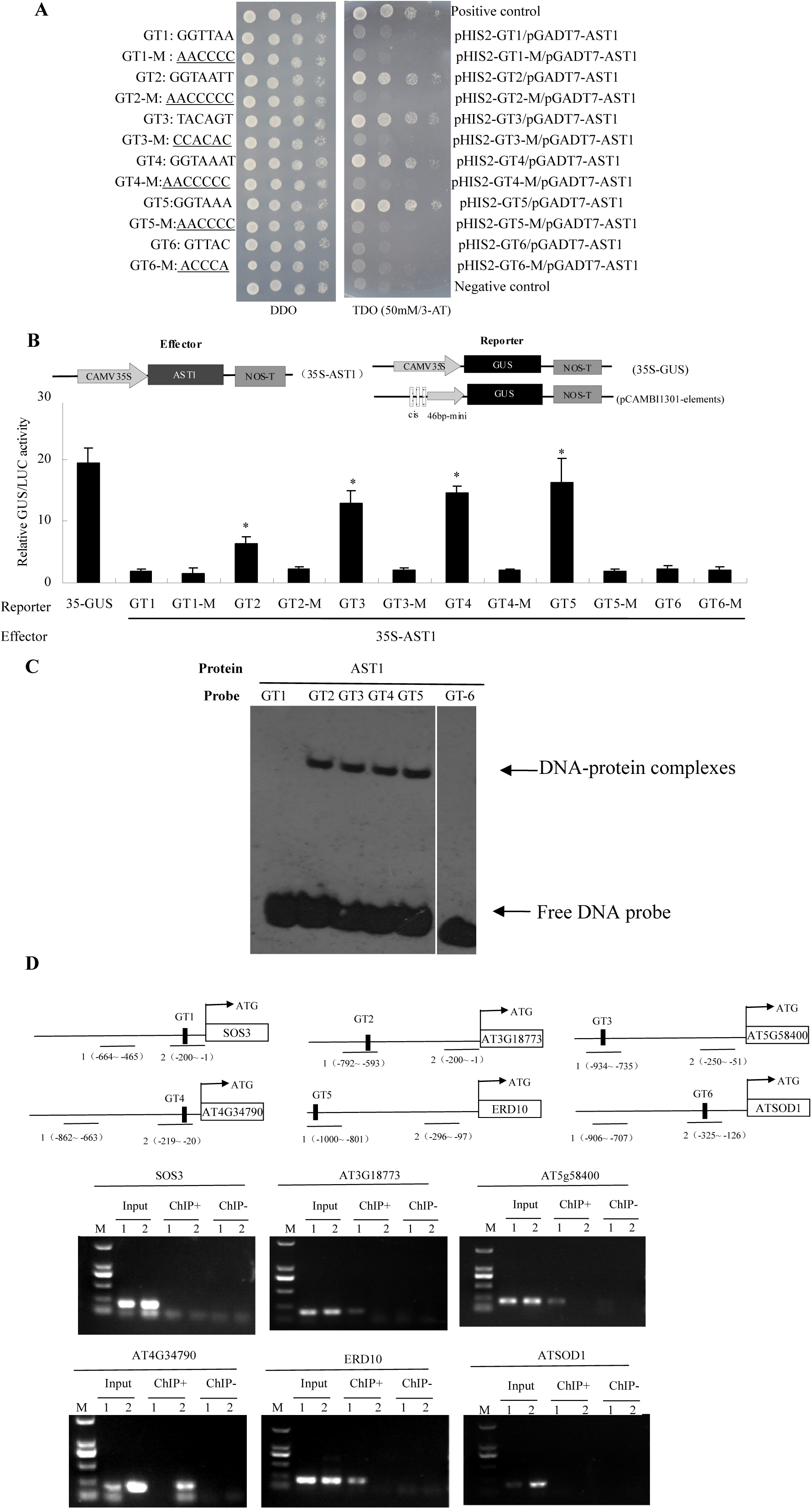
Identification of the GT motifs recognized by AST1. **(A)** Y1H assay of the GT elements recognized by AST1. Six GT elements and their mutations were respectively cloned in pHIS2 vector, and their bindings to AST1 were studied using Y1H. The above motifs were mutated following this principle, i.e. “A/T” was mutated to “C” and “C/G” was mutated to “A”. **(B)** Determination of the interaction of AST1 with GT motifs in tobacco plants. GT motifs and their mutations were fused separately with the 46-bp minimal promoter to drive *GUS* as reporters; each reporter was co-transformed with 35S:AST1 and 35S:LUC into tobacco. Diagrams of the reporter and effector vectors were shown. Data are means ± SD from three independent biological replicates. Asterisk (*) indicates a significant difference at P<0.05 compared with the mutations. **(C)** EMSA was carried out with AST1 protein and GT-box sequences. Lane 1-6, GT1, 2, 3, 4, 5 and 6 probes incubated with AST1 protein. The free probes and DNA-AST1 complexes were marked. **(D)** ChIP analysis of the binding of AST1 to GT elements. The promoters that contain only one type of GT motifs and no other motif recognized by AST1 were used in this experiment. Schematic diagram showing the positions of GT elements in the promoters. Input (input sample), ChIP+(immunoprecipitated with an anti-GFP antibody), ChIP-(immunoprecipitated with an anti-HA antibody).

Y1H results showed that AST1 could bind to this 12 base conserved sequences “AGAGAGAGAAAG” (Figure 6B). The 1st to 8th base of the 12 base conserved sequence appeared with the highest frequency and might be the core sequence of this motif, therefore, they were subjected for further study. The 1st to 8th base of 12 bp conserved sequences were represented by 32 types of sequences, and were all subjected to Y1H assays. The Y1H results showed that only some of the eight base sequences were bound by AST1; however, when the 7th base was G or the 8th was A, their binding to AST1 were lost (Figure 6B). Therefore, the eight base sequences that were bound by AST1 were represented by the consensus sequence [A/G][G/A][A/T]GAGAG, and was termed the AGAG-box. To further determine the bindings of the AGAG-box by AST1, we produced *GUS* gene reporter constructs that contained all the sequences of AGAG-box separately, the 12 base conserved sequence, or the six sequences (AGTGAGAA, AGTGAGGG, AGTGAGGA, GAAGAGAA, GAAGAGGG and GAAGAGGA) that could not be bound by AST1 according to Y1H (Figure 6B). Each reporter was co-transformed with the effector (35S:AST1) into tobacco plants. The GUS/LUC ratio showed that AST1 recognized all the AGAG-box sequences and the 12-base conserved sequence, but failed to bind to the other sequences (Figure 6C). This result was consistent with that of Y1H.

To further determine whether AGAG-box sequences could be bound by AST1, five types of AGAG-box sequences that showed highly transactivation when interacted with AST1 (Figure 6C) were labeled with biotin as the probes, and were used for EMSA. The results showed that the DNA-protein complexes were observed, and the complex binds were gradually decreased with increasing the unlabeled probes (Figure 6D), showing that AST1 could bind to these AGAG-box sequences. Meanwhile, the two sequences (AGTGAGGG and GAAGAGAA) that were not bound by AST1 according to Y1H were also studied, and EMSA result confirmed that they could not be bound by AST1 (Figure 6D).

To determine whether AST1 could bind to AGAG-box in *A. thaliana* plants, ChIP analysis was performed. Three genes whose promoters contained only AGAG-boxes and no GT motifs were used for ChIP analysis. The *ABR* gene (*AT3G02480*) whose promoter region did not contain both AGAG-box and GT motifs, and only contained an AST1 non-recognized sequence (GAAGAGAA) was used as the negative control. When using ChIP+ ( immunoprecipitated with the anti-GFP antibody) as the template, the promoter region containing the AGAG-box were PCR amplified; however, the promoter region far way from AGAG-box all failed PCR (Figure 6E), indicating that AST1 really bound to AGAG-box in *A. thaliana*. Additionally, the promoter region of *ABR* (containing GAAGAGAA that was not bound by AST1 according this study) also failed in PCR amplification when use ChIP+ as the template (Figure 6E). Meanwhile, the promoter regions could all be amplified from the Input, and the ChIP-( immunoprecipitated with the anti-HA antibody) failed PCR for all the promoter regions, indicating that ChIP results were reliable (Figure 6E). These results suggested that AST1 indeed bound to the AGAG-box to regulate the expression of genes in *A. thaliana*.

### AST1 binds to GT cis-acting elements

Previous studies showed that Trihelix proteins could bind to GT motifs, including GGTTAA (GT1), GGTAATT (GT2), TACAGT (GT3), GGTAAAT (GT4), GGTAAA (GT5) and GTTAC (GT6) (Green *et al*., 1987; Kay *et al*., 1989; O’Grady *et al*., 2001; Gao *et al*., 2009; Yoo *et al*., 2010). We first investigated the binding of AST1 to these GT motifs using Y1H. The results showed that AST1 bound to GGTAATT (GT2), TACAGT (GT3), GGTAAAT (GT4), GGTAAA (GT5), but failed to binds to GGTTAA (GT1) and GTTAC (GT6) (Figure 7A). The interaction between AST1 and these GT motifs were further performed in Tobacco. Three copies of each GT motif were fused with 35S minimal promoter to drive a *GUS* gene as a reporter, and were transformed with 35S:AST1 into Tobacco plants. The results showed that AST1 could bind to GT2, GT3, GT4 and GT5, but failed to bind to GT1 and GT6, which was consistent with the Y1H results (Figure 7B).

To further determine the bindings of AST1 to GT motifs, EMSA was performed. When the GT-1 and GT-6 probe was added, only the free DNA probe was observed, further indicating that GT1 and GT-6 were not bound by AST1. When GT2, 3, 4, and 5 sequences were respectively added with AST1 protein, the DNA-protein complexes could be observed (Figure 7C), confirming that GT2-5 sequences all could be bound by AST1.

To determine whether AST1 could bind to the GT motifs in *A. thaliana*, ChIP analysis was performed. Six genes whose promoters contained only GT1, GT2, GT3, GT4, GT5 or GT6 motifs, and no other known Trihelix binding motif (including the AGAG-box), were used for the ChIP analysis. When ChIP+ was used as the PCR template, the promoter fragments containing GT2, GT3, GT4, or GT5 motifs were amplified, and the promoter regions containing distant GT2, GT3, GT4, or GT5 motifs failed in PCR amplification. In addition, the promoter regions containing proximal or distal GT1 or GT6 motifs all failed in PCR amplification using ChIP+ (Figure 7D). At the same time, all the chosen promoter region could be amplified from the Input sample, and ChIP-failed to PCR amplify any of the promoter regions (Figure 7D), indicating that the ChIP-PCR results are reliable. These results together indicated that AST1 could bind to GT2, GT3, GT4 and GT5 (GT2-5), but not to GT1 and GT6 in *A. thaliana*.

### ChIP analysis of the genes directly regulated by AST1

To further determine the genes regulated directly by AST1, ChIP analysis was performed. The stress tolerance genes whose expressions were affected by AST1 according to qRT-PCR or RNA-seq were studied for ChIP analysis. The schematic diagram of the promoter fragments from different AST1-upregulated genes used for qChIP-PCR was shown as Figure S9. The results showed that besides *ABR, SOS3* and *ATSOD1*, the chosen promoter regions contained AGAG-box or GT2-5 motifs of all the studied genes were significantly enriched, suggesting that they were regulated directly by AST1 (Figure 8). Importantly, *SOS3, ABR* and *ATSOD1* did not contain an AGAG-box, GT2, GT3, GT4, or GT5 in their promoters, and their promoters could not be bound by AST1 (Figure 8). Furthermore, the expressions of *SOS3, ABR* and *ATSOD1* were not affected by AST1 according to the qRT-PCR results (Figure 4, Figure S6 and Figure S5b). These results further confirmed that AST1 could bind to the AGAG-box and the GT2, GT3, GT4, or GT5 (GT2-5) motifs to regulate the expression of genes. In addition, according to the ChIP results (Figure 8), the genes involving in water loss rate, ion homeostasis, and proline contents and ROS scavenging capability were mainly directly regulated by AST1.

**Figure 8.**
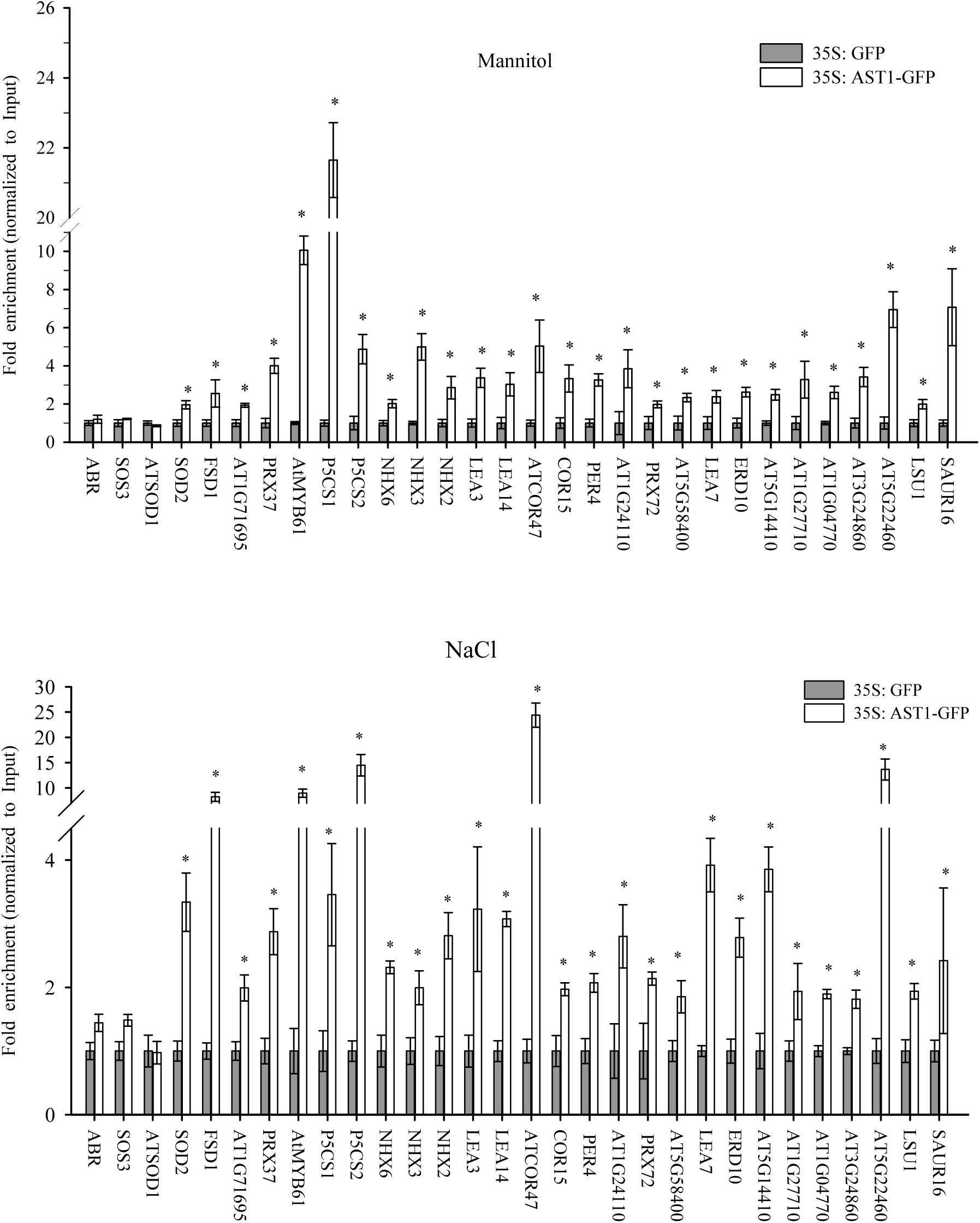
qChIP-PCR analysis of the genes directly regulated by AST1. Three-week-old 35S:GFP and 35S:AST1-GFP transgenic plants treated with 150 mM NaCl or 200 mM Mannitol were used for ChIP analysis. The promoter fragments that contained AGAG-box or GT elements identified by qRT-PCR and transcriptome were studied. The expression values in 35S:GFP plants were set as 1 to normalize the expression in 35S:AST1-GFP plants. *ABR, SOS3* and *ATSOD1* that were not regulated by AST1 and did not containing ASTA1 binding motifs were used as negative controls. *AT5G14410, AT1G27710, AT1G04770, AT3G24860, AT5G22460, LSU1, SAUR16* were the genes identified in RNA seq. The CDS of *ACTIN2*, which is not regulated by AST1, was used as internal control. Data are means ± SD from three independent biological replicates. Asterisk (*) indicates a significant difference at P<0.05 compared with the 35S:GFP.

## Discussion

AST1 is a GT transcription factor, whose function involved in abioic stress had not been characterized previously. In the present study, we identified the motifs bound by AST1 and further revealed the stress tolerance related genes regulated by AST1 and the physiological changes mediated by AST1 in response to abiotic stress.

### AST1 binds to a novel motif AGAG-box to regulate the expression of genes

Previous studies showed that some Trihelix proteins could bind to different types of GT-motifs (Kaplan-Levy *et al*., 2012). However, our results showed that AST1 could only bind to GGTAATT (GT2), TACAGT (GT3), GGTAAAT (GT4) and GGTAAA (GT5), but not to GGTTAA (GT1) and GTTAC (GT6) (Figure 7). Additionally, AST1 also binds to a novel motif, the AGAG-box, which contains eight types of sequences. Among these sequences, when the sequences “AAAGAGAG”, “AGAGAGAG”, “GGAGAGAG” and “GATGAGAG” were present, AST1 showed relatively higher activation of gene expression (Figure 6C). By contrast, when the other four sequences, “GAAGAGAG”, “GGTGAGAG”, “AATGAGAG” and “AGTGAGAG”, were present, AST1 showed relatively lower gene expression activation (Figure 6C). These results suggested that AST1 might show higher binding affinities to “AAAGAGAG”, “AGAGAGAG”, “GGAGAGAG” and “GATGAGAG” compared with those to “GAAGAGAG”, “GGTGAGAG”, “AATGAGAG” and “AGTGAGAG”. These two groups of sequences only had differences in the first to the third nucleic acids, indicating that these three nucleic acids might be relatively important for AST1 binding.

We screened the frequency of the occurrence of the AGAG-box and GT2-5 motifs in the promoters of genes regulated by AST1, including the 24 genes identified by qRT-PCR, and the 62 genes that were upregulated by AST1 according to RNA-Seq. Among these promoters, 58% (50 genes) contained AGAG-box motifs, and 65% (56 genes) contained different GT2-5 motifs. The occurrence frequencies of AGAG-box and GT2-5 motifs were similar, suggesting that like the GT motifs, the AGAG-box also played a very important role in AST1-mediated gene expression.

### AST1 binds to AGAG box and GT motifs serving as a transcriptional activator

We studied the binding of AST1 to different GT motifs or AGAG box in Tobacco plants. The results showed that AST1 could bind to AGAG box and GT2-5 to activate the expression of *GUS* gene (Figure 6C; Figure 7B), suggesting that AST1 should serve as a gene expression activator when binding to these motifs.

### The physiological response mediated by AST1

Plant guard cells form stomatal pores that played important roles in CO2 uptake for photosynthesis and in transpirational water loss. Transpiration accounts for most of the water loss in plants. Plants reduce transpirational water loss by inducing stomatal closure in response to drought stress (Munemasa *et al*., 2015). In the present study, we found that AST1 was highly expressed in guard cells (Figure 1A), and induces stomatal closure to reduce water loss (Figure 3). Previous studies showed that *AtMYB61* directly controls the stomatal aperture (Liang *et al*., 2005). Our study showed that AST1 could upregulate the expression of *AtMYB61* directly (Figure 3B and Figure 8). These results indicated that AST1 controlled stomatal closure and opening by regulating *AtMYB61* expression directly, thereby aiding water stress tolerance.

Maintenance of K^+^/Na^+^ homeostasis was quite important for plant salt tolerance (Sergey *et al*., 2007). Our study showed that AST1 reduced Na^+^ accumulation and decreases K^+^ loss (Figure 4). Meanwhile, AST1 also regulated genes involved in Na^+^ and K^+^ homeostasis, including *HKT1, NHX2, NHX3, NHX6* and *SOS2* (Figure 8).

These results indicated that AST1 could reduce Na^+^ accumulation and decrease K^+^ loss by regulating the expressions of Na^+^ and K^+^ transporter genes, which will contribute to alleviating salt stress.

Proline is the main solute used in osmotic potential adjustment. In *A. thaliana*, P5CS is the key enzyme in proline biosynthesis, and the degradation of proline is catalyzed by two enzymes, PRODH and P5CDH (Silva-Ortega *et al*., 2008; Szabados *et al*., 2010). Our results indicated that AST1 controlled the proline content and the expression of *P5CS* genes positively, and downregulates *PRODH* and *P5CDH* (Figure 5). These results suggested that AST1 induced the expression of *P5CS* to increase proline biosynthesis; simultaneously, it decreased the expression of *PRODH* and *P5CDH* to inhibit proline degradation, resulting proline accumulation to enhance osmotic potential, thereby improving salt and osmotic stress tolerance.

ROS scavenging is important for abiotic stress tolerance in plants. Excess ROS generated by abiotic stress attack all macromolecules, leading to serious damage to DNA, including lesions and mutations, cellular components, metabolic dysfunction and cell death (Karuppanapandian *et al*., 2011). Proline not only acts as osmotic adjuster but also serves as ROS scavenger. The proline content had been found to be highly induced by AST1 (Figure 5A). Additionally, SOD and POD are the two most important antioxidant enzymes in ROS scavenging. AST1 induced the expression of both *SOD* and *POD* genes to increase SOD and POD activities (Figure S5), which enhanced ROS scavenging capability and reduced ROS accumulation (Figure S5) to improve abiotic stress tolerance.

### AST1 regulates the expression of *LEA* genes to improve stress tolerance

The plant LEA family proteins, which are important for abiotic stress tolerance, stabilize the cell membrane, and serve as molecular chaperones or shield to prevent irreversible protein aggregation caused by abiotic stress, thus protecting the plant from damage (Serrano *et al*., 2003). Some LEA family proteins that had been confirmed to play a role in stress tolerance were studied here, and AST1 was found to induce the expression of most of the studied *LEA* genes (Figure S6 and Figure 8). Therefore, these *LEAs* highly expression would contribute to improve abiotic stress tolerance. Therefore, one of pathways that AST1 improved salt and drought tolerance was to induce the expression of *LEAs* involving abiotic stress tolerance.

In conclusion, our data suggested a working model for the function of AST1 in the abiotic stress response. Abiotic stresses, such as salt or osmotic stress, induce the expression of *AST1*. The induced AST1 protein binds to AGAG-boxes and/or GT2–5 motifs to regulate the expressions of genes involved in abiotic stress tolerance, such as stomatal aperture, K^+^/Na^+^ homeostasis, proline biosynthesis, ROS scavenging, and *LEAs*. The altered expressions of these genes lead to physiological changes, including reduced water loss and Na^+^ accumulation, prevention of K^+^ loss, elevated proline level, reduced ROS accumulation, and high expression of *LEAs*, which might play a role in stabilizing the cell membrane and serving as molecular chaperones to prevent protein aggregation caused by stress. These physiological changes ultimately improved abiotic stress tolerance (Figure 9).

**Figure 9.**
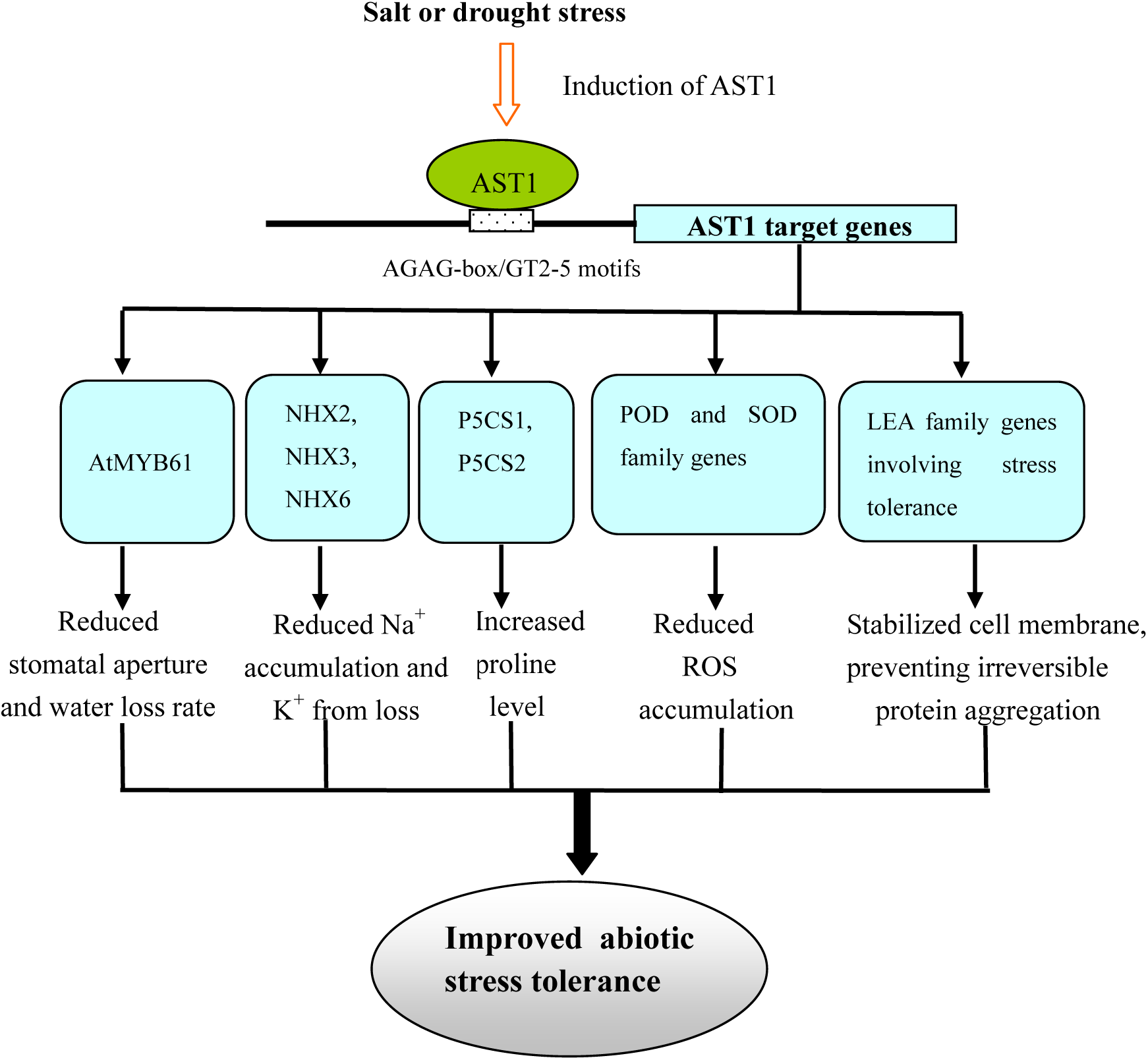
Working model of AST1 in response to abiotic stress. Abiotic stresses including salt or drought stress triggers the expression of *AST1*. Activated AST1 regulates the stress tolerance related genes by binding to the AGAG-box or GT2–5 motifs, which results in reducing stomatal aperture, water loss rate, Na^+^ accumulation, K^+^ loss, and ROS accumulation, increased proline level. The induced stress tolerance *LEA* genes may also play a role in stabilizing cell membrane and preventing irreversible protein aggregation. These physiological changes finally improve salt and drought stress tolerance.

## Acknowledgements

This work was supported by National Natural Science Foundation of China (No. 31270703), and the Fundamental Research Funds for the Central Universities (2572014AA25).

## Supplementary data

The following supplemental materials are available

**Supplemental Figure S1.** Subcellular localization of AST1.

**Supplemental Figure S2.** Relative expression of *AST1* in the OE and KO plants.

**Supplemental Figure S3.** Stress tolerance analysis on seedlings grown on 1/2 MS medium.

**Supplemental Figure S4.** Detection of cell death.

**Supplemental Figure S5.** Analysis of ROS levels and ROS scavenging capability.

**Supplemental Figure S6.** The expression of LEA family genes

**Supplemental Figure S7.** Hierarchical clustering analysis of the differentially regulated genes

**Supplemental Figure S8.** Go and KEGG analysis of differentially expressed genes.

**Supplemental Figure S9.** A schematic diagram of the promoter fragments used for qChIP-PCR.

**Supplemental Table S1.** Primers used in qRT_PCR

**Supplemental Table S2.** Primers used in Y1H

**Supplemental Table S3.** Primers used in tobacco Transient Expression Assay

**Supplemental Table S4.** Primers used in EMSA assay

**Supplemental Table S5.** Primers used in ChIP assay

**Supplemental Table S6.** Regualted genes by AST1 in RNA-seq

